# Exploring Conformational Transition of 2019 Novel Coronavirus Spike Glycoprotein Between Its Closed and Open States Using Molecular Dynamics Simulations

**DOI:** 10.1101/2020.04.17.047324

**Authors:** Mert Gur, Elhan Taka, Sema Zeynep Yilmaz, Ceren Kilinc, Umut Aktas, Mert Golcuk

## Abstract

Since its first recorded appearance in December 2019, a novel coronavirus (SARS-CoV-2) causing the disease COVID-19 has resulted in more than 2,000,000 infections and 128,000 deaths. Currently there is no proven treatment for COVID-19 and there is an urgent need for the development of vaccines and therapeutics. Coronavirus spike glycoproteins play a critical role in coronavirus entry into the host cells, as they provide host cell recognition and membrane fusion between virus and host cell. Thus, they emerged as popular and promising drug targets. Crystal structures of spike protein in its closed and open states were resolved very recently in March 2020. These structures comprise 77% of the sequence and provide almost the complete protein structure. Based on down and up positions of receptor binding domain (RBD), spike protein can be in a receptor inaccessible closed or receptor accessible open state, respectively. Starting from closed and open state crystal structures, and also 16 intermediate conformations, an extensive set of all-atom molecular dynamics (MD) simulations in the presence of explicit water and ions were performed. Simulations show that in its down position, RBD has significantly lower mobility compared to its up position; probably caused by the 6 interdomain salt bridges of RBD in down position compared to 3 in up position. Free energy landscapes based on MD simulations revealed a semi-open state located between closed and open states. Minimum energy pathway between down and up positions comprised a gradual salt bridge switching mechanism. Furthermore, although significantly lower than open state, ACE2 binding surface of RBD contained a partial solvent accessibility in its closed state.

## I. INTRODUCTION

In December 2019, a novel coronavirus named severe acute respiratory syndrome coronavirus 2 (SARS-CoV-2) causing the coronavirus disease (COVID-19) emerged as a human pathogen in the city of Wuhan, China. Since then, SARS-CoV-2 has spread worldwide rapidly. As of 17 April 2020, there are more than 2,000,000 confirmed COVID-19 cases globally, among them at least 135,000 have resulted in death (World Health Organization). Based on its genome sequence, SARS-CoV-2 belongs to same genus (betacoronavirus) as the severe acute respiratory syndrome coronavirus (SARS-CoV) and Middle-East respiratory syndrome coronavirus (MERS-CoV). SARS-CoV emerged in the Guangdong province of China in 2002 and resulted in an epidemic outbreak infecting 8,098 and killing 774 people (Drosten et al. 2003, Walls et al. 2020). MERS-CoV emerged in 2012 in the Arabian Peninsula, where it remains a major public health concern, having infected ~2519 and killed 858 people in 27 countries (Memish et al. 2020, Walls, Park et al. 2020). Phylogenetic analyses have suggested that SARS-CoV-2 shares 82% gene sequence identity with SARS-CoV (Lu et al. 2020, Wu et al. 2020, Zhu et al. 2020) and 51.8% gene sequence identity with MERS-CoV (Han and Yang 2020). Coronaviruses (CoVs) comprise a large and diverse family. Though having such high gene sequence identity with SARS-CoV and MERS-CoV, SARS-CoV-2 appears to be transmitted among humans more readily and much faster than SARS-CoV (Li et al. 2003, Wrapp et al. 2020) and MERS-CoV (Paraskevis et al. 2020, Zhu, Zhang et al. 2020). COVID-19 clinical symptoms are fever, dry cough, severe respiratory failure, and pneumonia (Wu, Zhao et al. 2020, Zhou et al. 2020). On 10 March 2020, COVID-19 was declared as a pandemic by the World Health Organization (WHO) (World Health Organization 2020). Currently there is no proven treatment for COVID-19. Thus, there is a pressing need for the development of vaccines and therapeutics for COVID-19.

Recognition of host cell and entry of the virus are the most critical steps determining the pathogenesis and viral infectivity in viral infections (Gallagher and Buchmeier 2001). Host cell recognition and fusion between host cell and viral membranes is provided by large trimeric spike (S) glycoproteins located on the CoV membrane (Fig. 1) (Belouzard et al. 2012). These S proteins interact with the human epithelial and respiratory cells via the angiotensin-converting enzyme 2 (ACE2) receptors of the host cells (Letko et al. 2020, Wrapp, Wang et al. 2020). Upon receptor binding and proteolytic cleavage, large-scale conformational changes in S protein structure occur and S protein interaction with the host cell membrane takes place facilitating the fusion process. Due to their critical role in the entry of coronaviruses into host cells, SARS-CoV-2 S proteins have emerged as a popular and promising therapeutic target. In the current study, structure and dynamics of the SARS-CoV-2 S protein is investigated via allatom molecular dynamics (MD) simulations to provide critical atomic level insight into the conformational transition mechanism of S protein from a receptor inaccessible closed to an accessible open state, which in turn has the potential to strongly benefit development of therapeutic strategies for COVID-19.

**FIG. 1.**
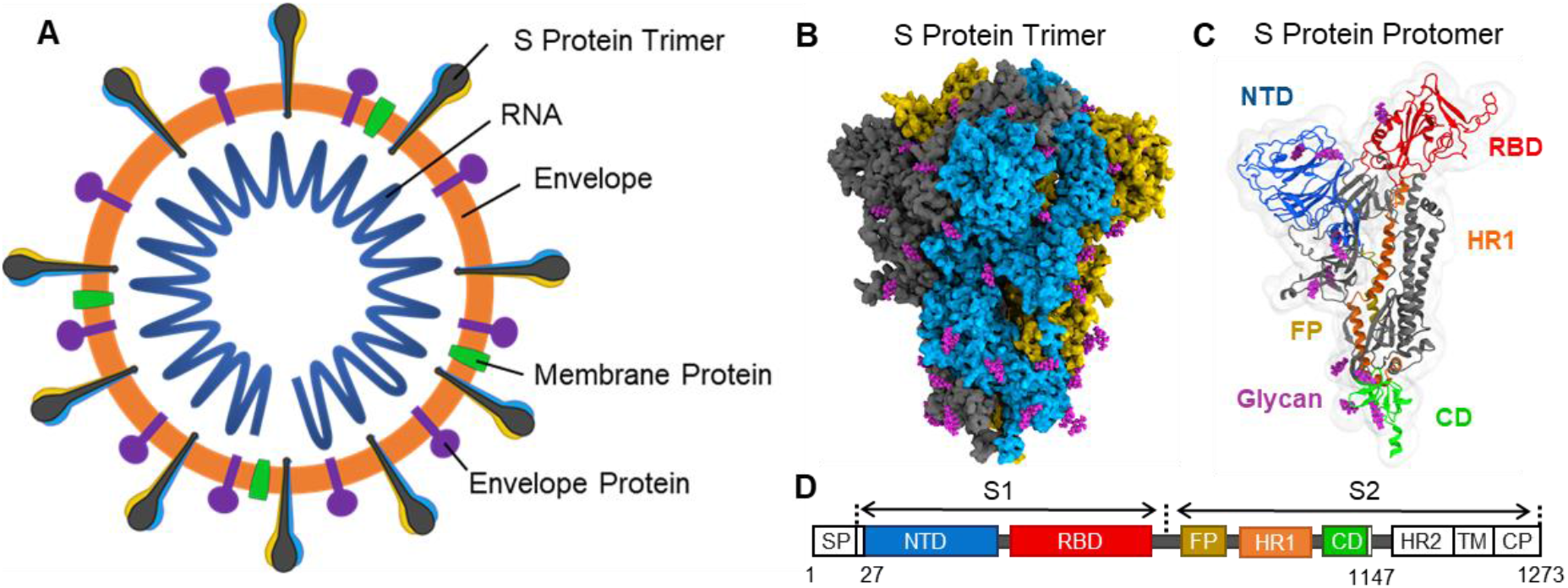
(a) Schematic representation of the structure of a coronavirus. (b) Crystal structure of the S protein (PDB ID: 6VXX). S protein is a large trimeric protein decorated with N-linked glycans (purple). S protein protomers, all of which are in down conformation, are shown in yellow, blue and gray. (c) S protein protomer structure in its down conformation is shown. Domains of an S protomer; NTD, RBD, FP, and HR1, CD, and glycans are shown in blue, red, mustard, orange, green, and purple, respectively. (d) The representative scheme of functional domains in S protein of SARS-CoV-2.

Each S protein protomer consists of two functional subunits (Fig. 1(c,d)): S1 subunit is responsible for binding to host receptor and S2 subunit contains the membrane fusion machinery (Walls et al. 2017, Kirchdoerfer et al. 2018). S1 subunit comprises two functional domains: N-terminal domain (NTD) and a receptor-binding domain (RBD), both of which are responsible for the binding to the host cell receptor (Li, Moore et al. 2003). S2 subunit contains three functional domains; fusion peptide (FP), heptad repeat (HR) 1, and HR2 (Jiang et al. 2020). Based on the RBD position, S protein can be in a receptor inaccessible closed or receptor accessible open state. As shown in Fig. 2, in the receptor inaccessible closed state, all RBDs are in the down position covering the S2 subunits; whereas in the receptor accessible open state, at least a single RBD is in an up position being rotated outwards from S2, hence exposing its receptor binding surface. Conformational switch of RBD from the down to up conformation is a prerequisite for ACE2 binding of the S protein. There are two cleavage sites in each protomer of the S protein. Among them, the S2’ site is located upstream from the FP. S1/S2 site, on the other hand, is located 103 residues downstream from S2’ and located at the boundary between the S1 and S2 subunits. Cleavage at S2’ sites is a requirement for fusion of host cell and virus membranes as it exposes the FPs at S2 to the environment and also separates the protein into S1-ectodomain and viral anchored S2 subunit. However, there is no consensus whether S1/S2 site cleavage is a requirement for S protein transition from pre to post fusion state. Upon S1/S2 site cleavage, S protein is divided into S1 and S2 subunits, which remain non-covalently bound in this metastable prefusion conformation (Bosch et al. 2003, Park et al. 2016, Kirchdoerfer, Wang et al. 2018). During cell infection, S protein binds to the host cell receptor ACE2 via RBD (Li, Moore et al. 2003, Wong et al. 2004, Zhou et al. 2020). Protein-receptor binding is expected to initiate a set of structural rearrangements in the S protein. Cleavage in S2’ promotes rearrangement of HR1 in the S2 subunit into an extended α-helix and insertion of the FP into the host membrane. Subsequently, a six-helix bundle comprising HR2 and HR1 is formed (Bullough et al. 1994, Bosch, van der Zee et al. 2003, Walls, Tortorici et al. 2017). This completes the transition of the S protein from its metastable pre-fusion state to a highly stable post-fusion state. S protein conformational transition results in the fusion of viral and host membranes, thus allowing the release of the viral genome into host cell (Walls, Tortorici et al. 2017).

**FIG. 2.**
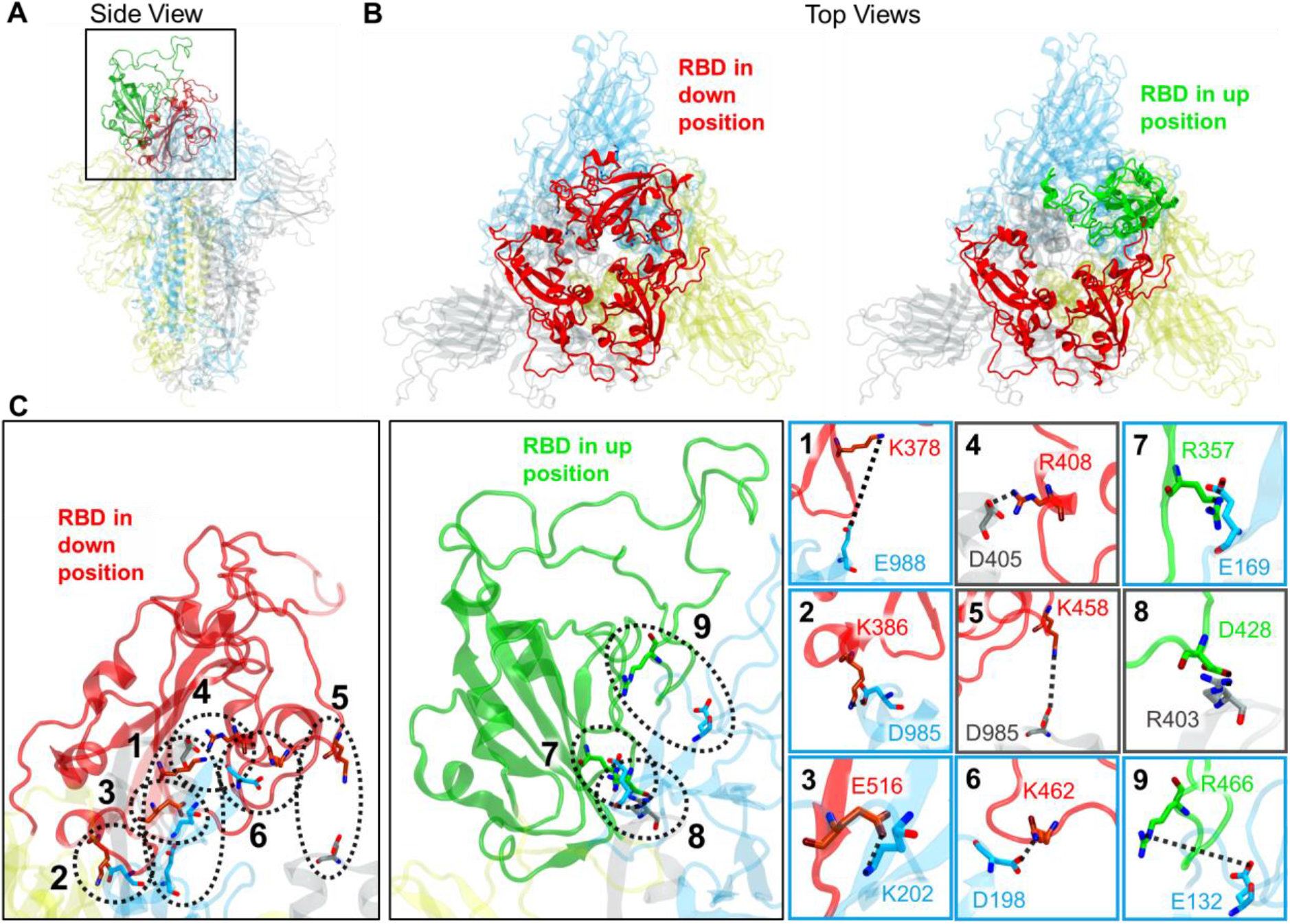
Down and up positions of RBD of S protein closed and open states. (a) The trimeric S protein conformation is shown from a side view. Closed and open state conformations sampled from MD simulations structures are aligned based on their secondary structures in S2 subunit. RBD of closed and open states are shown in red and green colors, respectively. Since the remaining structure is almost identical between up and down conformations, for the up conformation only the RBD is shown. Protomers A, B, and C are shown in gray, yellow, and light blue colors. (b) Closed and open states of S protein trimer is shown from top view. (c) Salt bridges observed between RBD and the neighboring protomers in closed (1 to 6) and open (7 to 9) states during MD simulations.

The first SARS-CoV-2 S protein structure was resolved in its open state and released on February 26 2020 with the PDB ID 6VSB. This structure had a 3.46 Å resolution and covers 76.8% of the sequence. Shortly after, on March 11 2020, SARS-CoV-2 S protein structures were resolved in their closed and open states having PDB IDs 6VXX and 6VYB. These closed and open structures have resolutions 2.8 Å and 3.2 Å, respectively, and each cover 76.4% of the protein sequence. Based on the crystal structures, the transition between the two states can be predicted to occur through a hinge-like conformational movement of RBD transiently hiding or exposing the determinants of receptor binding. As of today, there are no all-atom MD simulations studies published that investigated the dynamics of any of the abovementioned close-to-full length SARS-CoV-2 S protein structures (PDB IDs: 6VYB, 6VXX, and/or 6VSB). The earliest being on February 21 2020, a few preprints of computational studies were recently deposited to bioRxiv investigating binding between SARS-CoV-2 RBD and ACE2 (Peng et al. 2020, Shah et al. 2020, Smith and Smith 2020) and peptide/molecule binding to SARS-CoV-2 RBD (Zhang et al. 2020) via MD simulations. Upon performing MD simulations starting from the closed (PDB ID: 6VXX) and open (PDB ID: 6VYB) state structures, our current study shows that a salt bridge network between RBD and neighboring protomers exist in the closed state all of which is are gradually abrogated along the closed to open transition of the S protein. Furthermore, closed state is stabilized by newly formed salt bridges with the neighboring protomer. MD simulations showed that solvent accessibility of the ACE2 binding surface of RBD changes significantly upon the transition between open and closed states. Our simulations provided a description of the energy landscape for the transition between closed and open states of S protein by sampling the conformational space near the structurally resolved states as well as the region connecting these states. A new semi-open state was identified approximately midway along the transition pathway.

## II. MATERIALS AND METHODS

### A. Molecular dynamics system preparation

Coronavirus S protein structures having PDB IDs 6VYB (Walls, Park et al. 2020) and 6VXX (Walls, Park et al. 2020), which are in the closed and open states respectively, were used as starting structures in our MD simulations. As shown in Fig. 1, these crystal structures comprise most of the S protein domains; S1 subunit (NTD and RBD) and S2 (FP and HR1). Furthermore, S trimers are extensively decorated with N-linked glycans (Fig. 1), which are important for proper folding (Rossen et al. 1998) and for modulating accessibility to host proteases and neutralizing antibodies (Walls et al. 2016, Yang et al. 2016, Xiong et al. 2018, Walls et al. 2019). Closed and open state crystal structures of the S protein were obtained by performing the following mutation: (i) R682S, R683G, and R685G, which were performed to abrogate furin S1/S2 cleavage site, and (ii) K986P and V987P, which were performed for stabilization (stabilizing structure). The trimeric crystal structures comprise the sequence A27-S1147 of each protomer. However, out of these 1121 residues there are structural information missing for 149 of them in the closed state: V70-F79, Y144-N164, Q173-N185, R246-A262, V445-G446, L455-C488, G502, P621-S640, Q677-A688, L828-Q853. For the open state, on the other hand, there are 155, 172 and 161 residues missing for its three protomers (denoted by A, B and C) as follows: V70-N81, T114-Q115, Y144-N165, Q173-N185, A243-A262, S443-G447, E471-Y489, G502, P621-S640, Q677-S689, P812, L828-L854 of protomer A, A67-D80, L141-A163, Q173-N185, I197-G199, L212-R214, A243-A262, L455-L461, D467-F490, E516-P521, P621-S640, Q677-A688, P812, L828-Q853 of protomer B, and A67-D80, Y144-N165, Q173-N185, A243-A263, V445-G447, L455-L461, E471-F490, P621-S640, Q677-S689, P812, L828-F855 of protomer C. In order to model the missing residue stretches and add them to the protein structures, the three-dimensional structures of the closed and open states were modeled using SWISS-MODEL web server (Waterhouse et al. 2018) using 6VXX and 6VYB structures, respectively, as templates. For both closed and open states, the FASTA sequence spanning residues A27-S1147 for each S protein protomer is taken from NCBI having RefSeq: YP_009724390 (O’Leary et al. 2016). Prior to submission to SWISS-MODEL, all mutations in the crystal structures were reversed in the sequence. Using VMD (Humphrey et al. 1996), both closed and open crystal structures were separately solvated in water boxes each having at least 12 Å cushion of water in each direction. Ions were added to neutralize the system and NaCl concentrations were set to 0.150 M to represent a more typical biological environment. The size of the solvated S protein systems in the closed and open states were of ~410000 and ~415000 atoms in size, respectively.

### B. MD simulations

For each system, all-atom MD simulations were performed for an N, P, T ensemble in explicit solvent (water and ions) using NAMD 2.13 (Phillips et al. 2005) package with CHARMM36 (Best et al. 2012) force field. MD Simulations were performed at 310 K temperature and 1 bar pressure. Periodic boundary conditions were applied. Langevin dynamics was used to control the system temperature and pressure. All atoms were coupled to the heat bath. A time step of 2 fs was used. Two minimization-equilibration cycles were applied: All steps were applied under N, P, T conditions to relax water and find a local minimum of the whole system energy (Phillips et al. 2003). The energy of the initial system was first minimized for 10000 steps. Water was then equilibrated by keeping the protein fixed for 2 ns. Then, the system was minimized for 10000 steps. After the second minimization step, a harmonic constraint, which has 1 *kcal mol*^-1^ Å^-2^ spring constant, was applied to each Cα of the protein for 20 ns. At the third equilibration step, protein was released and the system was equilibrated for 5 ns. This equilibrium step is expected to be sufficient simulate the structural differences due to the radically different thermodynamic conditions of crystallization solutions and conventional MD simulations (Pullara et al. 2019). Finally, production MD simulations were performed. Atomic coordinates *R* of all atoms, pressures and the energies were recorded every 10 ps. Two sets of MD simulations were performed for each of the closed and open states of the S protein. A total length of 800 ns of MD simulations starting from the crystal structures were performed. Table I presents the complete list and length of each MD simulations performed in the present study.

**TABLE I.** Starting conformations and durations of the MD simulations performed.

| MD ID | Starting conformation | Simulation Type | Simulation Duration (ns) |
| --- | --- | --- | --- |
| 1a-b | Closed state crystal structure (6VXX) | MD | (a) 150, (b) 100 |
| 2a-b | Open state crystal structure (6VYB) | MD | (a) 450, (b) 100 |
| 3a-b | Closed state conformers (MD) | SMD | (a) 9, (b) 9 |
| 4a-b | Final conformers of MD 3a-b | MD | (a) 100, (b) 100 |
| 5a-b | Final conformers of MD 3a-b | SMD | (a) 2, (b) 2 |
| 6a-b | Final conformers of MD 5a-b | MD | (a) 100, (b) 60 |
| 7a-b | Final conformers of MD 5a-b | SMD | (a) 2, (b) 2 |
| 8a-b | Final conformers of MD 7a-b | MD | (a) 100, (b) 40 |
| 9a-b | Final conformers of MD 7a-b | SMD | (a) 2, (b) 2 |
| 10a-b | Final conformers of MD 9a-b | MD | (a) 100, (b) 80 |
| 11a-b | Final conformers of MD 9a-b | SMD | (a) 2, (b) 2 |
| 12a-b | Final conformers of MD 11a-b | MD | (a) 100, (b) 60 |
| 13a-b | Open state conformers (MD) | SMD | (a) 3, (b) 3 |
| 14a-b | Final conformers of MD 13a-b | MD | (a) 60, (b) 20 |
| 15a-b | Final conformers of MD 13a-b | SMD | (a) 2, (b) 2 |
| 16a-b | Final conformers of MD 15a-b | MD | (a) 20, (b) 20 |
| 17a-b | Final conformers of MD 15a-b | SMD | (a) 2, (b) 2 |
| 18a | Final conformers of MD 17a | MD | (a) 20 |

### C. Steered molecular dynamics

SMD (Isralewitz et al. 2001) is a MD simulation technique which allows to explore biological processes on time scales accessible to MD simulations. SMD has been successfully performed to examine a wide range of processes including molecule unbinding (Eskici and Gur 2013), domain motion (Izrailev et al. 1999), and protein unfolding (Lu et al. 1998). The main idea of SMD simulations is to pull the atom or atoms along a vector by applying external force. The pulling process can be carried out at a constant speed or by applying a constant force (Phillips, Isgro et al. 2003). Here, SMD simulations were performed using constant velocity; a dummy atom binds with a virtual spring to the center of mass of a group of atoms called steered atoms (SMD atoms) and is pulled along a selected pulling direction at a constant speed. The applied force depends on the instantaneous coordinates of the center of mass of the SMD atoms, **R** as follows (Phillips, Isgro et al. 2003),

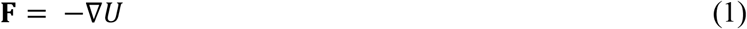

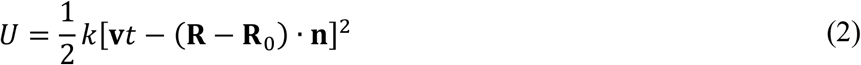

Here, *U* is the potential energy, **R**_0_ is the initial coordinates of the center of mass of SMD atoms, *k* is the spring constant, **v** is the pulling velocity, *t* is the time instant, and **n** is the direction of pulling.

The starting and end conformations of the SMD simulations were selected from our MD simulations of the closed and open states at time instances 20 ns and 40 ns. Cα atoms of RBD residues P337-A344, F347-V350, W353-S371, S375-Y380, V390-I410, G416-Y423, T430-S438, G447-R454, L492-Q498, Q506-L517, T523-G526 of protomer B were selected as SMD atoms, since they show minimal deformation upon superposition of up and down RBD conformations. In the first 20 ns of closed state MD simulations residues K986, D985 of protomer A and K113, D985, E988 of protomer C interacted with RBD of protomer B. Thus, their Cα atoms were kept fixed during our SMD simulations. Pulling vector was constructed by first aligning closed and open state conformations via S2 of protomer B and subsequently constructing a vector pointing from the center of SMD atoms in the closed state conformation to the open state conformation. A pulling velocity of 1 Å/ns was selected and spring constant is selected as 50 *kcal/mol*/Å^2^. These conditions satisfy stiff-spring approximation (i.e. dummy atom followed the center of mass of SMD atoms closely), while the spring constant was soft enough to allow small deviations (Isralewitz, Gao et al. 2001).

### D. Principle component analysis

Protomer conformations sampled in MD simulations were aligned with protomer B of the open state crystal structure using the C_α_ atoms of the S2 domain helices and beta sheets (Fig. S1). Upon alignment, the covariance matrix **C** was constructed using the 3×459 dimensional configuration vector **R** composed of the instantaneous C_α_ atom coordinates of the RBD and S2 domain helices and beta sheets as follows,

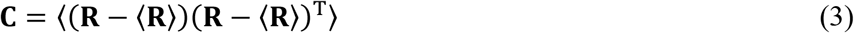

Here, 〈**R**〉 is the trajectory average of **R**. Principal components (PCs) are obtained by performing eigenvalue decomposition of **C** as,

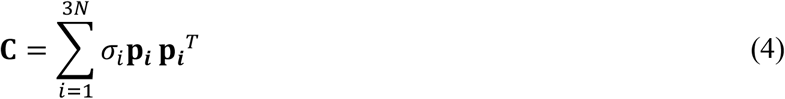

Here, **p_i_** is the *i*^th^ PC and *σ_i_* is the corresponding variance. Thus, *σ_i_* scales with the magnitude of motion along **p**,. PCs are ordered in descending order with respect to their *σ_i_* values. **p_i_** (or PC1) has the largest variance, **σ**_1_ thus represents the most dominant motion. Similarly, PC2 is the second most dominant motion observed in the MD trajectory.

## III. RESULTS AND DISCUSSION

### A. RBD in closed state shows less mobility than in open state, likely due to a higher number of interdomain salt bridges

Starting from the closed and open crystal structures, two sets of MD simulations were performed for each of the closed and open states. As shown in Fig. 2, in the open state only a single RBD is in up position whereas for the closed state all 3 RBDs are in down position. In accord with the crystal structure, protomers in the open state will be denoted with A, B, and C, among which protomer B is in up conformation. Superposition of protomers in the closed state indicate that all protomers have almost identical structures, showing root mean square deviation (RMSD) values of ~1 Å based on the helix and beta sheet Cα atoms. Thus, each MD simulation for the closed state will generate 3 MD simulation trajectories of the down conformation of the protomer. In order to investigate the mobility of the RBD domain in the closed and open states of the S protein trimer, the distance between RBD and S2 domains were tracked throughout the MD trajectories. As shown in Fig. 3, the distribution of the distance between RBD and S2 demonstrated a significantly wider distribution in the RBD up position compared to that of the RBD down position (in the closed state MD simulation). Standard deviations of the RBD-S2 distance distributions were 1.3 Å for the up RBD and 0.5 Å for the down RBD. Thus, RBD in up conformation in the open state clearly shows higher mobility than RBD in down conformation in the closed state.

**FIG. 3.**
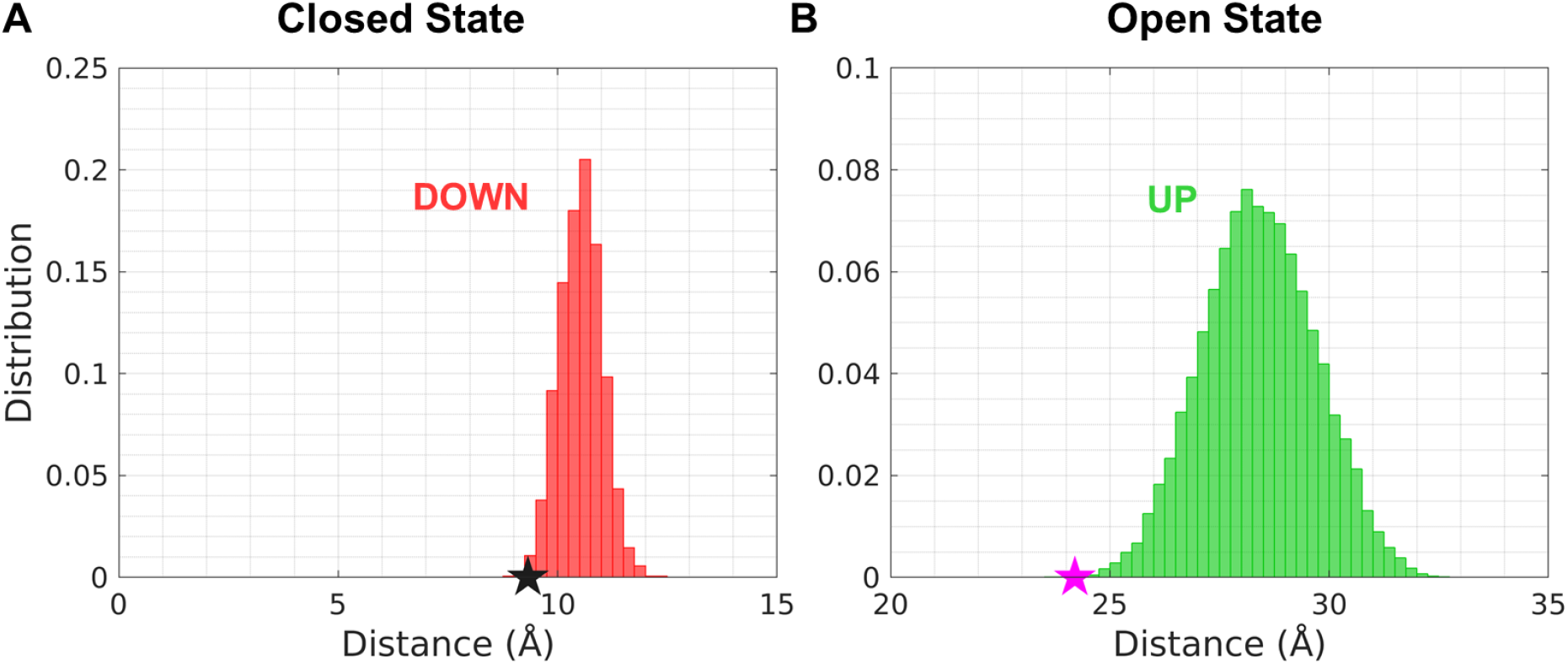
Distribution of the distance between RBD and S2 domain observed in MD simulations. The normalized distribution of the interdomain distances between RBD and S2 domain is shown for protomer B in (a) the closed state and (b) open state. The interdomain distance was calculated by evaluating the average of the distance between the C_α_ atoms of residue pairs K378(RBD)-E988(S2) and K386(RBD)-D985(S2). Distances in crystal structures are marked with black (PDB ID: 6VXX) and magenta (PBD ID: 6VYB) stars.

Superposition of the RBD structures in the down and up position via the Cα atoms of the helices and beta sheets shows a minimal structural difference of 0.5 Å RMSD. Furthermore, RBD domain conformations were conserved throughout our MD simulations. Thus, based on the crystal structures and our MD simulations it can be predicted that RBD undergoes a rigid body motion. Since there is no noticeable structural difference between the RBD structures, the difference in the RBD mobility in the open and closed state is expected to be caused by differences in interdomain interactions of RBD. To this aim, interdomain salt bridges of RBD with the remaining parts of protein were investigated in the closed and open states. Based the closed state crystal structure, RBD forms the following three interdomain salt bridges with residues of neighboring protomer; K378-E988 and K386-D985 with domain S2, and E516-K202 with NTD. Significantly, upon performing MD simulations in the closed state the following additional interdomain salt bridges of RBD formed with its neighboring protomers: R408-D405 with RBD, K458-D985 with S2 domain, and K462-D198 with NTD (Fig. 2). The distribution of the distances between the basic nitrogens and acidic oxygens of these 6 interdomain salt bridges of RBD are shown in Fig. 4. As can be seen the most prevalent salt bridge observed in MD simulations was R408-D405. Compared to its down position, RBD made significantly less interdomain salt bridges in its up position. Although there were no interdomains salt bridges present for RBD of protomer B in its up conformation in the open state crystal structure, MD simulations demonstrated the formation of the following interdomain salt bridges of RBD of protomer B with its neighbors: D428-R403 with RBD of protomer A, and R357-E169 and R466-E132 with NTD of protomer C. The distributions of the distances between the basic nitrogens and acidic oxygens of these 3 salt bridges are depicted in Fig. 4(b).

**FIG. 4.**
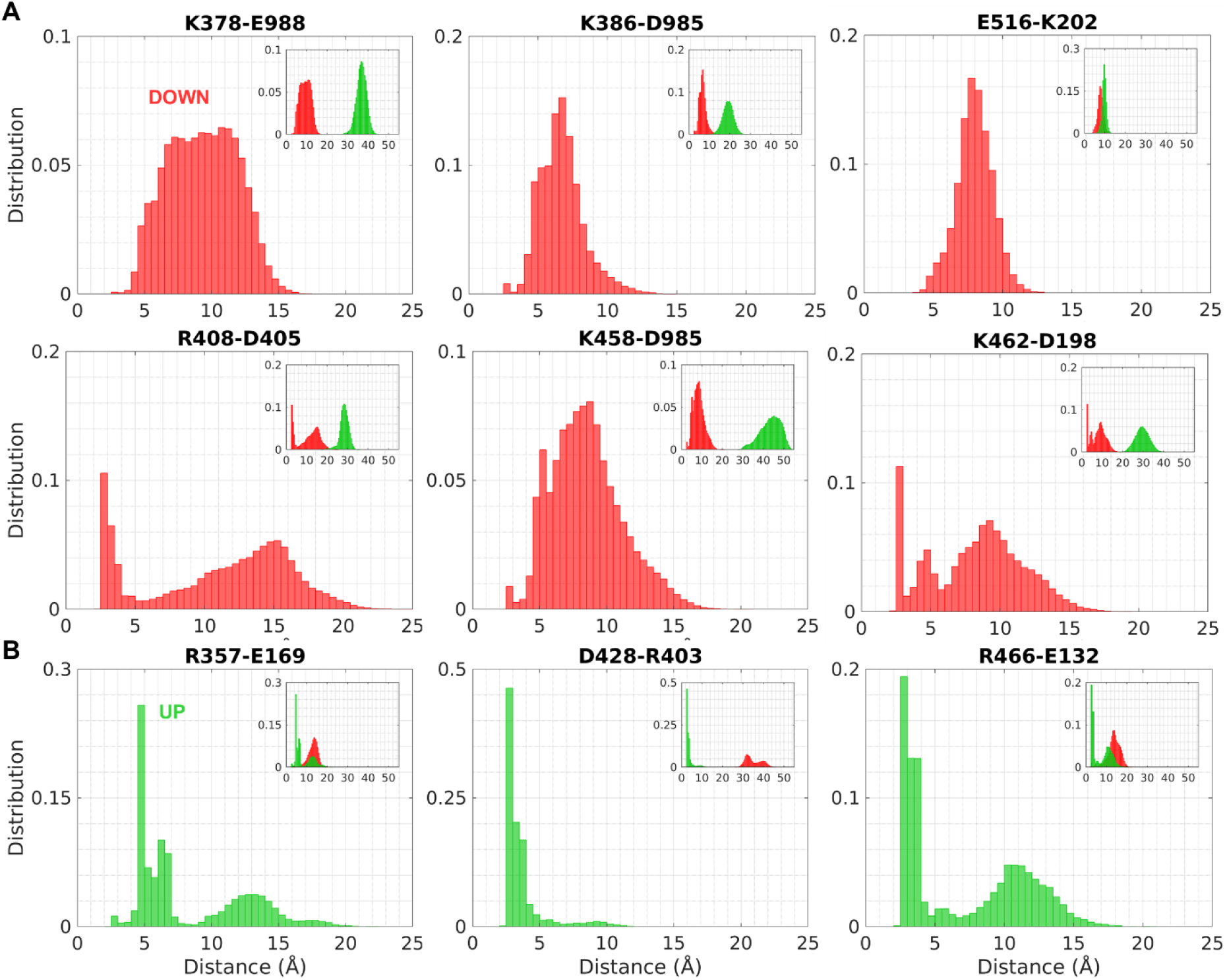
Distribution of the distances between basic nitrogens and acidic oxygens of interdomain salt bridges of RBD during MD simulations. Distribution of distances between basic nitrogens and acidic oxygens of the indicated amino acids are provided for MD simulations in the (a) closed state and (b) open state. For a pair of amino acids forming salt bridges, the first index indicates the RBD amino acid whereas the second one indicates the amino acid on neighboring protomers. The insets show the comparison for each salt bridge in closed and open states.

### B. Solvent accessibility of the ACE2 binding surface of RBD is significantly higher in the open state

Recently, the crystal structure of RBD bound to ACE2 was resolved at 2.45 Å resolution (PDB ID: 6M0J (Lan et al. 2020)). Based on this structure, RBD amino acids K417, G446, Y449, Y453, L455, F456, A475, F486, N487, Y489, Q493, G496, Q498, T500, N501, G502, Y505 (locations shown in Fig. 5) are interacting with ACE2. In order to investigate the solvent accessibility of these amino acids, close contact water molecules were calculated for the closed and open state MD simulations. To this aim, water molecules within 5 Å of the RBD amino acids involved in ACE2 binding were evaluated for each protein conformation sampled in the MD simulations. Notably, while the number of close contact water molecules were 433 for RBD in up position on average, the number of close contact water molecules was only 296 for RBD in down position of the closed state. As shown in Fig. 5(a), the RBDs of the S protein trimer are closely packed in the closed state, hence strongly limiting water accessibility to amino acids K417, Y453, L455, F456, and Y505. The remaining amino acids G446, Y449, A475, F486, N487, Y489, Q493, G496, Q498, T500, N501, and G502 are solvent accessible and could potentially be accessed by small molecules in the closed state of S protein. Furthermore, superposition of RBD bound ACE2 structure onto the closed state S protein trimer shows a large steric clash between ACE2 and RBD (Fig. 5(b)). Thus, making it spatially impossible for ACE2 to bind RBD in closed state.

**FIG. 5.**
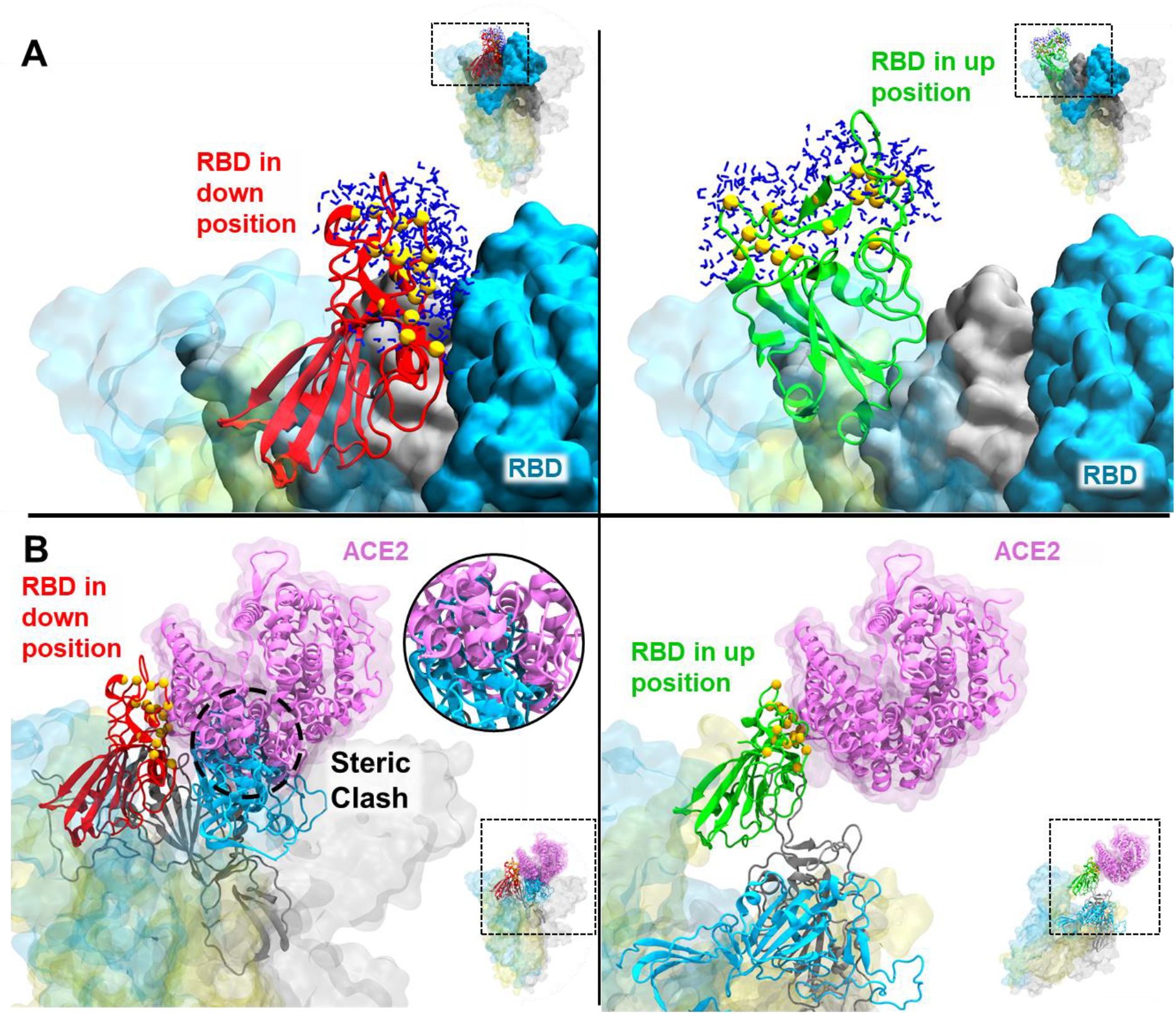
Solvent and ACE2 accessibility of ACE2 binding surface of RBD. (a) Close contact water molecules at the ACE2 binding surface of RBD of protomer B are shown for the closed state (left panel) and open state (right panel) in dark blue colored licorice representation. (b) ACE2 binding pose is shown on RBD in the down and up conformation of protomer B. Steric clash between ACE2 and RBD is highlighted in a circle. RBD amino acids involved in ACE2 interactions are highlighted with yellow beads.

### C. Energy landscape based on MD simulations of (starting from) closed and open S protein trimeric structures demonstrates well defined down and up states for S protein protomers

In order to explore the most dominant features of the protein dynamics in the closed and open states, PCA was performed using all conformations of protomer A, B and C sampled in the closed state MD simulations and all protomer B conformations sampled in the open state MD simulations. PCA is an effective and proven method used to dissect the most prominent motions of a protein along a given MD trajectory (Please refer to Sec. II D for details) (Gur et al. 2018). For a MD trajectory of a system comprised of N atoms, PCA provides 3xN modes of motions, among which PC1 and PC2 represent the first and second most prominent motion observed in the MD trajectory. PC1 and PC2 obtained from the combined MD trajectory of down conformation of protomer in closed state and up conformation of protomer in open state is shown in Fig. 6(a). The ratio of the variances between PC1 and PC2 are *σ*_1_/*σ*_2_ = 69.35, and cumulatively they account for 98.6% of the total motion in the MD trajectory; i.e. the first two PCs account for 98.6% of the total variance. As can be seen in Fig. 6(a), PC1 describes a rigid body like motion of RBD. PC1 alone is able to describe 98% of the structural transition between the protomer in its down an up conformation (Loop regions and NTD were not included in this calculation). PC2 is also characterized by predominant movement of RBD.

**FIG. 6.**
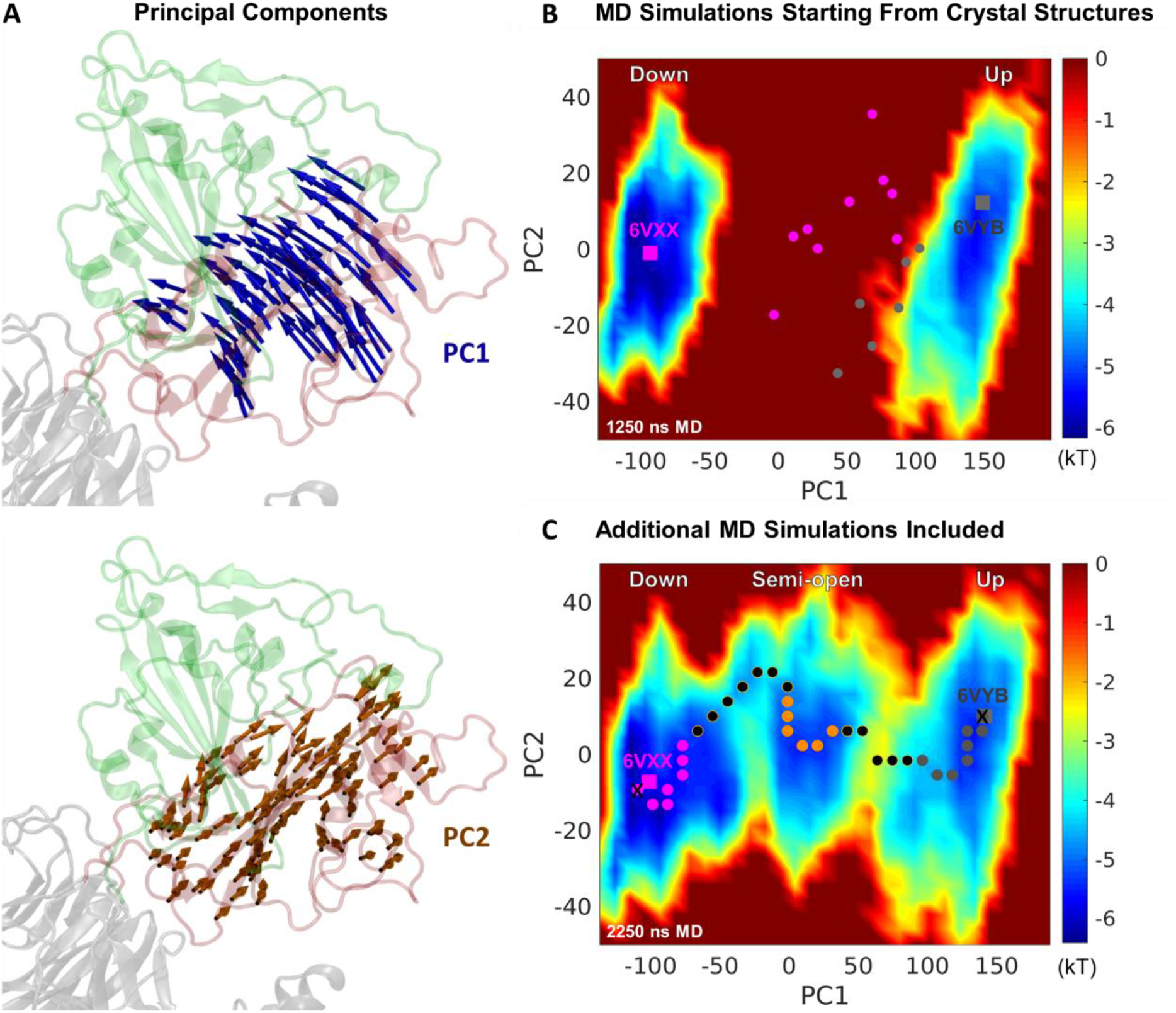
The first two principal components and the energy landscapes obtained from MD simulations. (a) The first two principal components of the MD simulations data superimposed on protomer B structure. (b) Energy landscape for S protein protomers obtained using closed and open state MD simulations (totaling 1250 ns). Reaction coordinates defining x and y axis were defined as PC1 and PC2 obtained upon combination of closed and open state MD simulations. The closed (6VXX) and open (6VYB) crystal structures are shown with magenta and gray squares. The initial conformers for MD simulations, which are obtained from up-to-down and down-to-up SMD simulations are indicated with gray and magenta dots, respectively. (c) Energy landscape for S protein conformers obtained using all MD simulations (total 2250 ns). Minimum free energy pathway is shown with dots. Magenta, orange, and gray dots represent the part of the pathway sampling the down, semi-open and up states. End point of minimum energy pathway is shown with crosses.

Conformations sampled in MD simulations were projected onto PC1 and PC2. Each PC was divided into 30 bins, hence generating an 30×30 grid system. Subsequently, based on the occupancy of these grids, distributions of the projections *f*(**R**) were computed along sets of PCs. Lower negative values along PC1 indicate an increased level of RBD closure, whereas higher positive values represent increased degree of RBD opening. Using distributions along PC1 and PC2, free energy surfaces were calculated as, *A*(**R**) = —*kTln* (*f*(**R**)) + constant (Gur et al. 2013). The free energy surface of the S protein protomer projected on PC1 and PC2 is shown in Fig. 6(b). The free energy surface shows two distinctive energy wells, representative of down (on the left) and up (on the right) conformational states of S protein protomer. MD simulations starting from closed state resulted in a narrower distribution along PC2, while distribution widths along PC1 was comparable. The region between down and up conformational states was not sampled by the MD simulations, probably due to the inability of MD simulations to simulate global transitions between these two states in the absence of any bias.

### D. New semi-open state emerges upon performing MD simulations starting from intermediate conformations between down and up states of S protein protomer

In order to explore the region between the down and up states on the energy surface and determine the minimum free energy pathway connecting these states, we performed a new round of unbiased MD simulations starting from unpopulated regions between the down and up states. To this aim, SMD simulations were performed starting from closed state conformations in order to provide intermediate conformations from which new set of unbiased MD simulations will be initiated. SMD simulations were performed by steering RBD of protomer B towards its up position, which will be denoted with down➔up. Similarly, SMD simulations starting from open state conformations were performed pulling RBD of protomer B towards its down position, which will be denoted with up➔down. Each of down➔up and up➔down SMD simulations were performed by starting from 2 separate starting conformers sampled from the closed and open state MD simulations, respectively. Starting from S protein conformations sampled from SMD simulations at 9 Å, 11 Å, 13 Å, 15 Å, and 17 Å along the down➔up SMD pulling direction (Table I, MD IDs 3, 5, 7, 9, and 11), and 3 Å, 5 Å and 7 Å along the up➔down SMD pulling direction (Table I, MD IDs 13, 15 and 17), a new set of 16 unbiased MD simulations were performed (Table I, MD IDs 4, 6, 8, 10, 12, 14, 16, 18). Combining the protomer B trajectories of the new set of 1000 ns long unbiased MD simulations with our first set of trajectories provided a total of 2250 ns of S protein protomer data to be used for our analysis. All protomer B conformations sampled in unbiased MD simulations were projected onto PC1 and PC2, which were obtained in the first round of MD simulations (MD1-2, Table I). Distribution of projections were computed and the free energy surface shown in Fig. 6(b,c) was obtained. Free energy surface revealed a new semi-open state located halfway across the transition between the down and up states, and a few additional substates at various locations. The semi-open intermediate state is separated from the down and up states by energy barriers of ~3kT. The energy barrier connecting the down and semi-open states provided a passage significantly lowering the energy requirement to move across this barrier. For the energy barrier between the semi-open and up states, on the other hand, three small passages, none of which is significantly lowering the energy requirement to pass over the energy barrier, were observed.

### E. Transition between down and up states comprises a switching mechanism of salt bridges

A minimum free energy pathway connecting the down and up states were constructed upon following low energy regions on the energy landscape while gradually proceeding towards the target state. The location of the conformations, which were identified to be on the minimum free energy path, are shown in Fig. 6(c). The pathway circumvents the energy barrier between the down and semi-open states by passing through the passage. For the transition between the semi-open and up states the mild passage in the middle of the energy barrier was selected as the crossing point since it was the shortest path connecting the low energy regions of the semi-open and up states. In order to have an understanding of the behavior and contribution of salt bridge breakage and formation during the conformational transition between the down and up state, the transition pathway was divided into 32 bins and salt bridges of all conformations located inside each bin were investigated. For each bin the distribution of the distance between the acidic oxygen and basic nitrogens of amino acids pairs K378-E988, K386-D985, E516-K202, R408-D405, K458-D985, K462-D198, R357-E169, D428-R403, and R466-E132 were evaluated. The bin averages of distances are depicted in Fig. 7. It is to be noted that standard deviations of distances are not shown in Fig. 7. As can be seen in distributions of Fig. 4, for the listed interactions, the salt bridge forming requirement of acidic oxygen and basic nitrogens to be closer than 6 Å is not satisfied throughout the trajectory, yet strong interactions are conserved even at slightly higher distance. Thus, if conformers inside a bin are forming salt bridges frequently and otherwise strong interactions are preserved for the amino acid pairs, we will refer to this bin as a salt bridge forming bin for the sake of conciseness of the discussion.

**FIG. 7.**
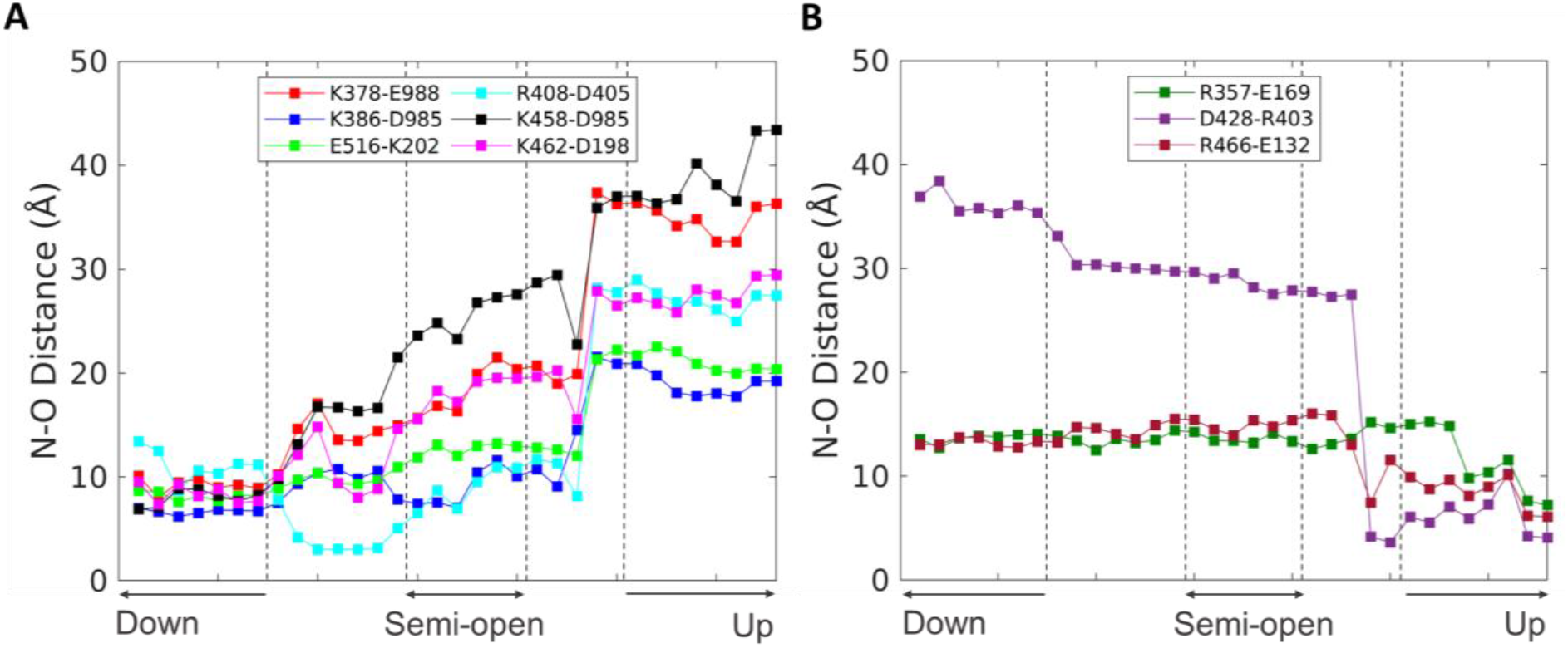
Time evolution of salt bridges during the transition between the down and up conformations. Distance changes between the basic nitrogens and acidic oxygens throughout MD simulations for the salt bridges observed in the (a) down and (b) up states.

As can be seen from the distance distributions in Fig. 7, during the closed to semi-open transition, the salt bridge R408-D405 with RBD of protomer A is formed, and K458-D985 with S2 of chain A, K378-E988 with S2 of protomer C and K462-D198 with NTD of protomer C are broken. The closed to semi-open transition takes place by passing through a substate, which differentiates from the down state by the formation of R408-D405 and breakage of K378-E988 and K458-D985. This switch in salt bridges is probably the reason why this substate is observed along the transition between the closed and semi-open state. Transition from this substate into the semi-open state comprises breakage of K462-D198 with NTD of protomer C. Significantly, crossing the energy barrier, which separates the semi-open state from the up state, to settle into the up state requires the breakage of 3 salt bridges; K386-D985 with S2 of protomer C, R408-D405 with RBD of protomer A, and E516-K202 of NTD of protomer C; and formation of two salt bridges; R466-E132 with NTD of protomer C and D428-R403 with RBD of protomer A. Salt bridge R357-E169 with NTD of protomer C is formed later on along the transition pathway, probably further stabilizing the up state.

## IV. CONCLUSION

MD simulations of the closed and open states show that the RBD in its up position shows higher mobility than RBD in its down position in the closed state. Analysis of the interdomain salt bridges of RBD suggest that the reason behind the higher mobility is the significantly lower number of interdomain salt bridges that RBD is forming in its up position. Interestingly, although solvent accessibility of the ACE2 binding interface of RBD is significantly limited in the closed state, there is still a significant amount accessible to the solvent. In fact, out of 17 amino acids of the ACE2 binding interface in RBD, 12 are solvent accessible in the closed state. This is a significant finding, as it potentially implicates that small molecule binding to inhibit the S protein in its closed state could be theoretically possible. Superposition of the ACE2 bound RBD structures clearly shows that it would be impossible for the S protein to bind ACE2 in its closed state. Thus, in contrast to an inhibitor binding to the open state, an inhibitor binding to the closed state would not need to compete with ACE2 for the binding interface. Energy landscapes based on our MD simulations suggest the existence of an intermediate semi-open state for the S protein, in which RBD of one protomer is halfway between its down and up positions. This semi-open state shows a distinct salt bridge network from the down and up states, and it remains to be explored whether this semi-open state could potentially be an effective drug binding target for therapeutics. Taken all together, our study provides unique atomic-level insight into the dynamics, interactions, and the solvent accessibility of the ACE2 binding surface of the S protein in its closed and open states. Moreover, conformational details and energetics of the down to up transition, which effectively puts the S protein in an active form, were explored. This extensive novel insight regarding S protein structure, dynamics, and energetics is expected to support the global drug and vaccine development efforts.

## ACKNOWLEDGEMENTS

We gratefully acknowledge the support of the National Center of High Performance Computing (UHeM) at ITU, Turkey. We thank Dr. Kadir Diri for his technical support at UHeM.

